# Physical interactions reduce the power of natural selection in growing yeast colonies

**DOI:** 10.1101/332700

**Authors:** Andrea Giometto, David R Nelson, Andrew W Murray

## Abstract

Microbial populations often assemble in dense populations in which proliferating individuals exert mechanical forces on the nearby cells. Here, we use yeast strains whose doubling times depend differently on temperature to show that physical interactions among cells affect the competition between different genotypes in growing yeast colonies. Our experiments demonstrate that these physical interactions have two related effects: they cause the prolonged survival of slower-growing strains at the actively-growing frontier of the colony and cause faster-growing strains to increase their frequency more slowly than expected in the absence of physical interactions. These effects also promote the survival of slower-growing strains and the maintenance of genetic diversity in colonies grown in time-varying environments. A continuum model inspired by overdamped hydrodynamics reproduces the experiments and predicts that the strength of natural selection depends on the width of the actively-growing layer at the colony frontier. We verify these predictions experimentally. The reduced power of natural selection observed here may favor the maintenance of drug-resistant cells in microbial populations and could explain the apparent neutrality of inter-clone competition within tumors.

**Significance Statement:** Microbes often live in dense populations such as colonies and biofilms. We show that the success and extinction of yeast strains within a growing colony are determined by a combination of their relative fitness and the forces exerted by proliferating cells on their neighbors. These physical interactions prolong the survival of less-fit strains at the growing frontier of the colony and slow down the colony’s takeover by fitter strains. This reduction in the power of natural selection favors the maintenance of genetic diversity in environments in which the strains’ relative growth rates vary with time. Growth-induced physical interactions may thus favor the maintenance of drug-resistant cells, which are typically less-fit than non-resistant cells, within dense microbial populations.

## Introduction

Life in the microbial world often occurs in crowded environments, such as colonies, biofilms, and tissues. Within such dense populations, cells compete for nutrients and space to survive, proliferate, and propagate their genotype to future generations. Classical results of population genetics theory on the competition between different genotypes in well-mixed populations (1) do not apply directly to these microbial aggregates, due to their growing size and spatial structure (2-9). Microbial range expansion experiments are a valuable tool for investigating ecological and evolutionary dynamics in spatial contexts (4, 5, 10-15) and extend our understanding of spatial population genetics (16). Early studies of microbial range expansions have shown that the growth of microbial colonies causes the spatial separation of genotypes (2), which makes theoretical models derived for well-mixed populations inapplicable. Biological interactions between genotypes can either reduce this segregation or increase it, depending on the nature of the interactions. For instance, Müller et al. (10) showed that mutualistic yeast strains cannot de-mix if they are obligate mutualists and thus depend on each other for growth. In contrast, McNally et al. (17) showed that antagonistic interactions among bacterial strains cause a phase separation that further increases the spatial separation of different strains. Although additional deviations from classical theory could be produced by the mechanical interaction between cells in dense populations, the effects of such interactions on microbial competition is still relatively unexplored. Among the few exceptions, Farrell et al. (18) showed that the physical properties of a cell can affect the dynamics of beneficial mutations in growing microbial colonies. They showed that the probability that a mutant spreads in the population can be related to summary statistics characterizing cell alignment and the roughness of the expanding front. In parallel to our investigation, Kayser et al. (19) showed that the mechanical interaction between a faster-growing and a slower-growing strain can cause the prolonged survival of the slower-growing one at the colony frontier, a process that we have investigated in a different geometry and from a different modeling perspective.

In this work, we study the effect of mechanical interactions among cells on the competition of strains in growing colonies of the budding yeast, *Saccharomyces cerevisiae*. We focus on the competition for space and for nutrients of two yeast strains with different fitnesses and study the dynamics of selective sweeps (the fitter strain’s increase in relative frequency at the frontier of the expanding colony) and the local extinction dynamics of the less-fit strain. In *S. cerevisiae* colonies, growth occurs mostly in close proximity to the outer boundary, and so we consider a strain to be extinct if it lags behind during the expansion and thus exits the growth zone. We find that mechanical interactions reduce the strength of natural selection by slowing down the extinction of slower-growing strains and by reducing the rate at which faster-growing ones increase their frequency within a population. We show that a continuum model inspired by overdamped hydrodynamics can reproduce the experimental data and make new predictions that we verify experimentally. Specifically, the model predicts that the speed at which selective sweeps spread in a growing colony is inversely related to the width of the actively growing layer at the colony frontier. We tested the model by co-culturing two yeast strains with different temperature-dependent growth profiles. We show that the prolonged survival of the less-fit strain, which occurs via the formation of thin, persistent filaments (Fig. 1A-A2), promotes the maintenance of genetic diversity in the population when two strains compete in an environment in which their relative growth rates vary with time. This reduction in the efficiency of natural selection may be beneficial for microbial populations by delaying the extinction of slower-growing genotypes which carry traits that can later be beneficial for the population, such as resistance to antimicrobial drugs.

**Fig. 1.**
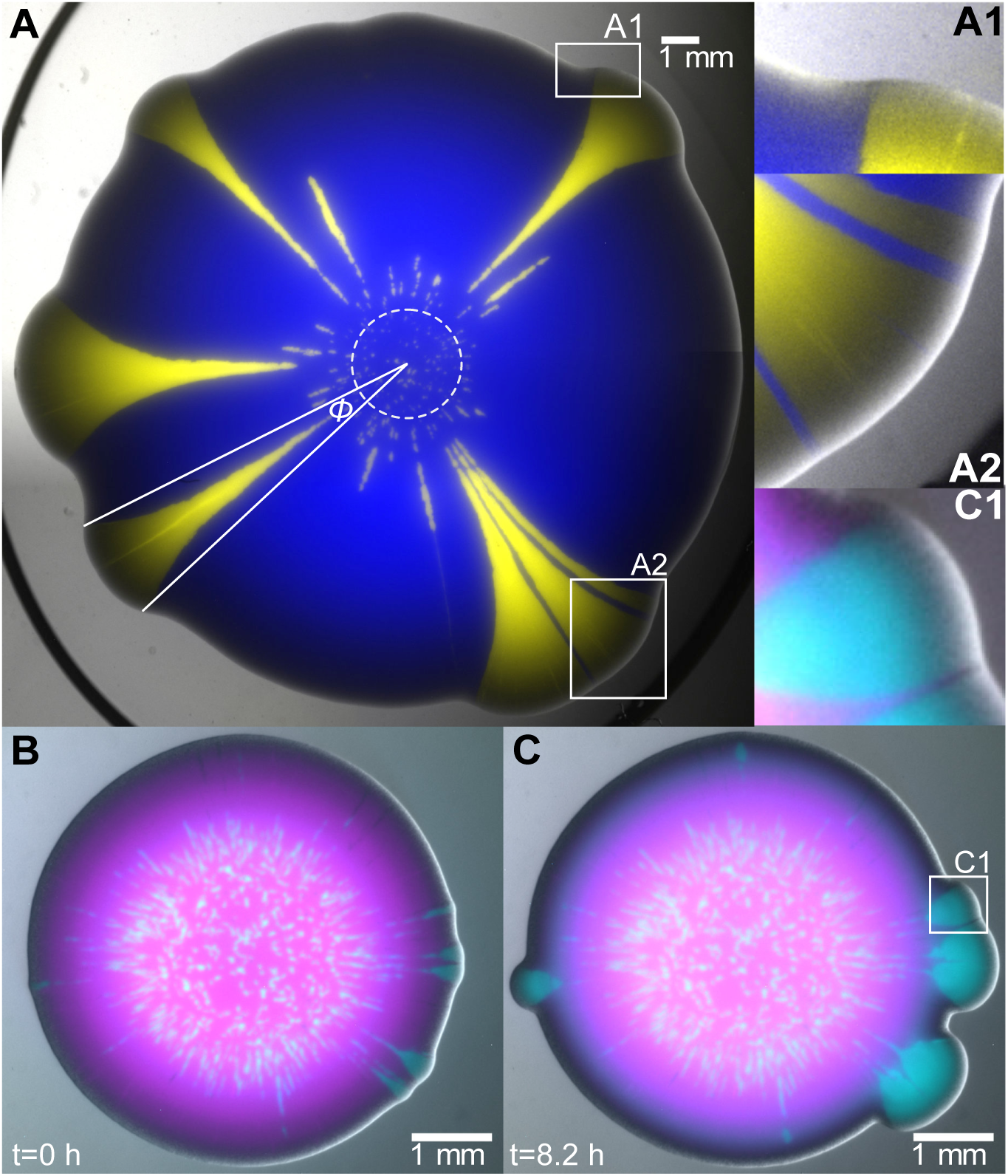
*S. cerevisiae* colony-growth experiments hint at physical interactions between strains. (A) Combined fluorescent and bright-field image of a colony, four days after the inoculation of a mixed droplet of strains yAG1 (blue) and yAG2 (yellow) at relative frequencies of 99% and 1%, grown at 30°C. The white dashed circle marks the inoculum size. Because yAG2 divides faster at 30°C, some single-strain yellow sectors expand their opening angle *ϕ* as the colony grows radially, while others go extinct due to stochastic effects in the early stages of growth. By expanding faster, the faster-growing strain displaces the other strain at both sides of yellow sectors (A1) and can enhance the expansion speed of the slower-growing strain when it is trapped between two yellow sectors (blue filaments in A2) and is carried along. (B-C) Even non-proliferating cells (magenta) can be pulled outwards by a proliferating (cyan) strain. The images show combined fluorescent and bright-field images of the same colony at early and later times. In this experiment, the non-heat-sensitive strain yAG19 (cyan, initial frequency 10%) and the heat-sensitive strain yAG20 (magenta, initial frequency 90%) were grown at 28°C for two days and subsequently imaged with a stereoscope incubated at 37°C, a temperature at which yAG20 cells cannot divide. Panel B shows the first snapshot at which the heat-sensitive strain stopped expanding (Fig. S1D). Panel C shows the colony after a further 8.2 h. (C1) Close-up of the region marked with a white square in C, highlighting the displacement of the non-growing, heat-sensitive strain next to expanding sectors of the growing strain and an expanding magenta filament trapped between two cyan sectors. Because the heat-sensitive strain did not divide between B and C, its displacement is due to the physical interaction between the two strains.

## Results

When two yeast strains are mixed and inoculated as a drop on a solid nutrient medium, they form a mixed-strain colony that expands radially with time. A well-known feature of such colony expansion is the spatial de-mixing of strains due to genetic drift (2), which forms single-strain sectors. When one strain expands faster than the other, the size of its sectors increases with time (5) (Fig. 1A). We performed colony-growth experiments using two *S. cerevisiae* strains with different doubling times. As expected, single-strain colonies of the strain (yAG2) with the larger growth rate (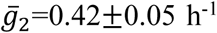, median±SD) expanded faster than single-strain colonies of the strain (yAG1) with the smaller growth rate (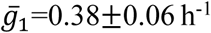, median±SD). We found that single-strain sectors within mixed colonies interact mechanically in ways that affect the competition between the two strains. These mechanical interactions are visible in Fig. 1A. At each side of sweeping sectors of the faster-growing (yellow) strain, the slower-growing (blue) strain was displaced and pulled towards the exterior of the colony (Fig. 1A-A1). Occasionally, the slower-growing strain was trapped between two sectors of the faster-growing one in the early phases of the expansion and traveled longer distances (blue filaments in Fig. 1A-A2) than in other regions of the colony, where the slower-growing strain was farther from the faster-growing strain. The enhanced expansion of the slower-growing strain occurs as a thin, persistent filament (composed of several thousand cells, not to be confused with bacterial filamentous cells) that originates via a combination of cell division and the physical interactions with the faster-growing strain, the most prominent of which is the pressure exerted by proliferating cells on their neighbors.

We wanted to rule out the possibility that persistent filaments were due to the ability of the faster-growing strain to speed up (for example, via chemical secretions) the proliferation of cells of the slower-growing strain that were nearby. We therefore produced mixed colonies from two strains: a wild-type strain that proliferated at temperatures from 20 to 37°C (yAG19, cyan in Fig. 1B-C) and a strain carrying a temperature-sensitive mutation that could only proliferate at temperatures below 30°C (yAG20, magenta in Fig. 1B-C). To ask if the formation of persistent filaments required the proliferation of the less-fit strain, we imposed the extreme condition of preventing the proliferation of the less fit strain. The heat-sensitive strain used here is a *cdc26*Δ mutant that arrests in mitosis and therefore cannot complete the cell division cycle at temperatures above 30°C (20). We grew single-strain and mixed-strain colonies of these two strains at 28°C for two days and subsequently imaged them with a stereoscope incubated at 37°C for 23 h. We observed that single-strain, heat-sensitive colonies stopped expanding after 9 h from the start of the measurement (Figs. 1B and S1). In the following hours, single-strain sectors of the non-heat-sensitive strain emerged from the mixed colony, displacing the heat-sensitive strain and producing patterns (Fig. 1C-C1) that are visually similar to those shown in Fig. 1A. Specifically, the non-dividing heat-sensitive strain was nevertheless displaced on each side of sweeping sectors of the non-heat-sensitive one and sectors of the heat-sensitive strain trapped between two sectors of the other strain (Figs. 1C1, S2 and Movie S1) were squeezed and formed filaments that resemble those shown in Fig. 1A-A2. Because the heat-sensitive single-strain colonies were not expanding when non-heat-sensitive sectors emerged from the mixed-strain colony (Fig. S1D), the displacement of the heat-sensitive strain was caused by the physical interaction between the two strains and the formation of thin filaments did not require the proliferation of the less fit strain.

The experiments described above show that mechanical interactions cause the prolonged survival of the slower-growing strain in mixed-strain colonies, via thin, persistent filaments trapped between sweeping sectors of the faster-growing strain. To highlight the effect of this persistence on the competitive dynamics of strains in growing colonies, we designed a colony-growth experiment in which we could use environmental switches to alternate which strains grew faster and slower. The strain yAG2 is cold-sensitive due to the *trp1-1* mutation, which requires it to import tryptophan from the medium, a process whose efficiency is reduced at lower temperatures (21), so that yAG1 grows faster than yAG2 at low temperatures and more slowly at high temperatures. At 12°C, single-strain colonies of yAG1 expanded faster (0.288±0.004 mm/d) than those of yAG2 (0.226±0.010 mm/d), whereas at 30°C single-strain colonies of yAG2 expanded faster (1.15±0.02 mm/d) than those of yAG1 (0.96±0.03 mm/d). Thus, by changing the temperature during a mixed-strain colony growth experiment, we could control which strain was proliferating faster. We performed mixed-strain colony growth experiments starting from a 50%-50% mixture of yAG1 and yAG2 and incubated these colonies at 30°C at the start of the experiment, then moved the colonies to 12°C after a few days, and moved them again to 30°C towards the end of the experiment (Methods). The duration of each temperature phase varied between different replicates. Fig. 2 and Figs. S3-S5 show that strains can survive as persistent, thin filaments at the temperature where they grow slower and then expand in the next phase, at a temperature where they grow faster than their competitor.

**Fig. 2.**
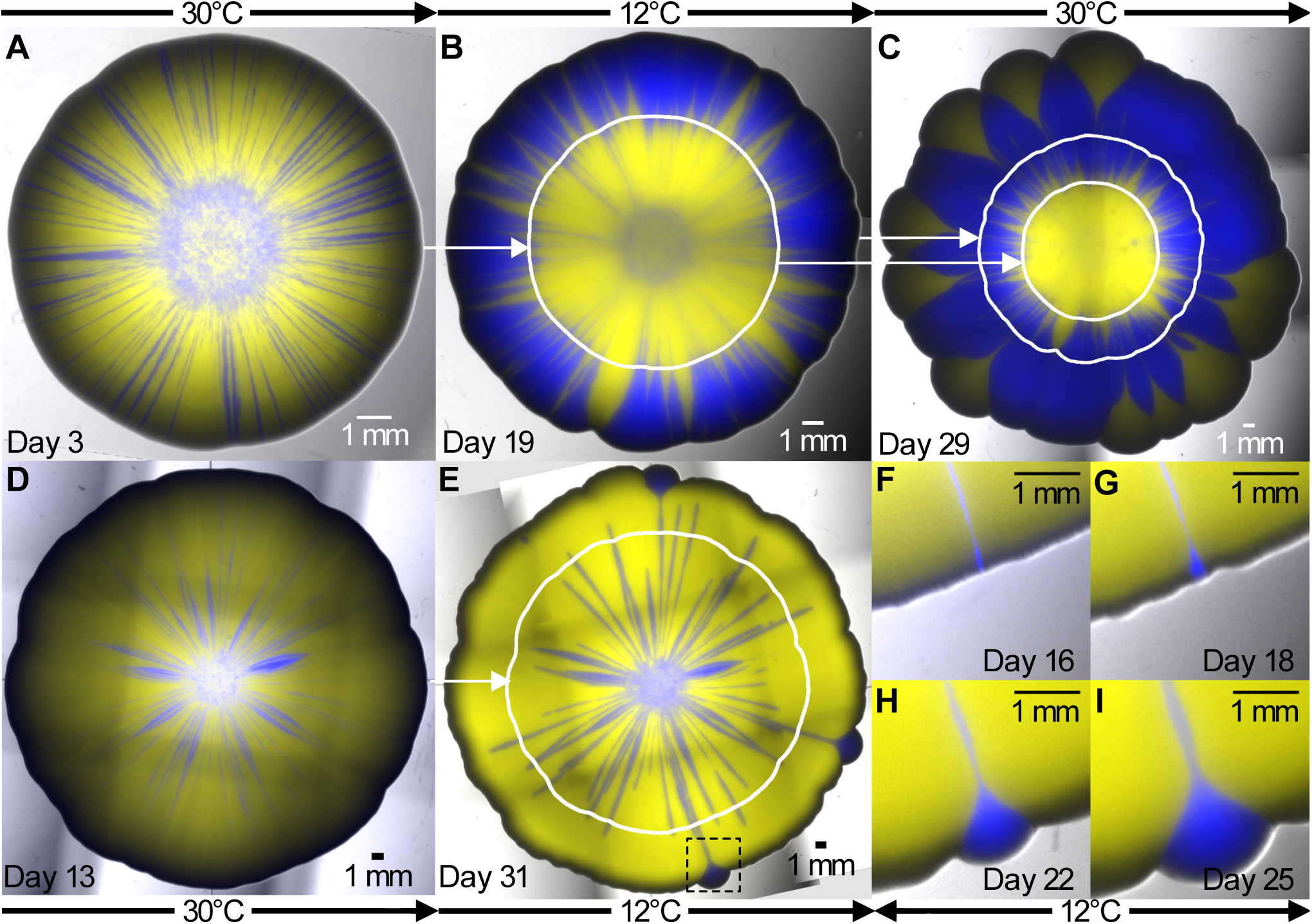
Colony-growth experiments in time-varying environments highlight the relevance of mechanical interactions to the competition between strains in mixed colonies. (A) Mixed yAG1 (blue, 50% initial frequency, the faster-growing strain at 12°C) and yAG2 (yellow, 50% initial frequency, the faster-growing strain at 30°C) colony grown at 30°C for 3 days. The colony was then grown for 16 days at 12°C (B) and for further 10 days at 30°C (C). (D) A different, mixed yAG1 (blue, 50% initial frequency) and yAG2 (yellow, 50% initial frequency) colony grown at 30°C for 13 days. The colony was then grown for 18 days at 12°C (E). At each temperature, the slower-growing strain survives as thin, persistent filaments that allow it to recover at the colony frontier following temperature changes. White curves show the colony perimeter at the previous temperature switch. Pictures were taken immediately before temperature changes (A, B and D) and at the end of the experiments (C and E). Panels F-I show the recovery of the strain yAG1 (in the region corresponding to the black, dashed square in E) from a thin filament during growth at 12°C. Temperatures shown indicate the incubation temperature during the days immediately preceding the images. The colony sizes are rescaled to be approximately equal in radius.

We made a hydrodynamic model to describe the observed interactions between strains at the macroscopic scale (i.e., at scales much larger than the typical cell diameter). Our model for the dynamics of the competition between two strains in growing colonies used a two-dimensional continuum description based on overdamped hydrodynamics. This approach is valid on time scales that are long compared to a cell division time and length scales that are large compared to a cell diameter and to the colony height near the frontier. We first describe the model for a single-strain colony and then generalize it to the case of multiple strains. Let *ρ*(*x,y,t*) be the two-dimensional density (with dimensions of cells/length^2^) of a single-strain colony at time *t*, where *x* and *y* are cartesian coordinates on the agar plane. If *h* is the height of the colony, we have *ρ* = *hρ*_3*d*_, where we assume that the three-dimensional density *ρ*_3*d*_ (with dimensions of cells/length^3^) is constant within the colony. Colony expansion in the vertical direction is much slower than the expansion in the horizontal plane of the agar and thus experimental colonies are approximately flat in the direction parallel to the agar surface. We therefore assume that *h* (and thus *ρ*) is also constant within the colony, and equal to zero outside. Let *g*(*x,y,t*) be the local growth rate (with dimensions of 1/time) of yeast cells in position (*x,y*) at time *t*. Experimental observations reveal that the two-dimensional spatial distribution of strains in mixed-strain colonies is approximately time-invariant at distances larger than 500 μm (about 100 cell diameters) behind the colony frontier (SI section 9), indicating that cell divisions that affect the visible surface of the colony take place mostly within this distance from the frontier. This observation led us to define a distance *δ* from the frontier of the colony such that 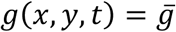, with constant 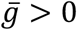, for every point (*x,y*) inside the colony within a distance *δ* from the frontier at time *t*, and *g*(*x,y,t*) = 0 for every other point. From a hydrodynamic perspective, cell growth is seen as a source of fluid with space and time-dependent rate *g*. With these assumptions, the radial expansion speed of a circular single-strain colony is equal to 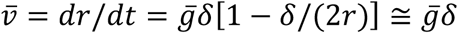 (SI Methods), where the last approximation is justified because typically *δ* ≪ *r* in colony-growth experiments. The considerations made above translate into the following equation for the two-dimensional velocity field 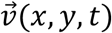 which displaces cells within the colony:

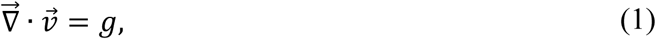

where 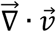 is the divergence of 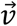. Eq. 1 enforces mass conservation within the colony along a streamline, ensuring that the two-dimensional density *ρ* is constant inside the colony. Within the colony, the two-dimensional pressure field *p*(*x,y,t*) satisfies the equation:

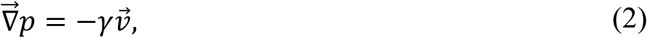

which states that the velocity 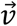 is proportional to the pressure gradient 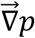 and inversely proportional to *γ*, the frictional drag applied to the colony by the substrate. Eqs. 1 and 2 have been used previously to model biofilm growth (22). We show in the SI section 3 that they can be derived from the three-dimensional continuity and Navier-Stokes equations by adopting the lubrication approximation (23), which gives the value of the friction coefficient *γ* = 2*μ*/*h*^2^ in terms of the colony dynamic viscosity *μ* and the colony height *h*. The approximation is appropriate when 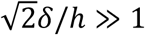, that is, when the height of the colony is small compared with the growth layer width, as found experimentally (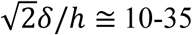, depending on the experimental setup). Upon taking the divergence of Eq. 2 and using Eq. 1, one finds:

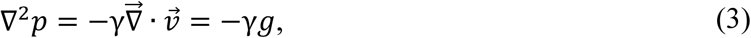

that is, the pressure field within the colony satisfies a Poisson equation, with the boundary condition *p* = *p*_0_ at the outer edge of the colony, where *p*_0_ is a constant reference pressure (because the value of *p*_0_ does not affect the dynamics, we set *p*_0_ = 0). This simple model can be generalized to mixed-strain colonies by considering two different two-dimensional densities *ρ*_1_ (*x y t*) and *ρ* (*x y t*), giving the density of each strain. Given the sharp boundaries between the two strains observed in colony growth experiments (see, e.g., Fig. 1A), we assume that the two strains cannot simultaneously occupy the same position in space at the same time *t*, so that for each point (*x y*) within the colony either *ρ*_1_ or *ρ*_2_ must be equal to zero, and the colony density is equal to *ρ* = *ρ*_1_ + *ρ*_2_. In the same way, the local growth rate is equal to *g* = *g*_1_ + *g*_2_ within the colony, where either *g*_1_ or *g*_2_ must be equal to zero, according to which strain occupies the position (*x y*) at time *t*. With this definition of *g*, the velocity field and pressure in the mixed-strain model are still given by Eqs. 2 and 3. Note that, because pressure is proportional to *γ* (Eq. 3) and the velocity field is given by 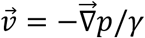 (Eq. 2), the frictional coefficient *γ* drops out and its value does not affect the displacement of strains within the colony and thus its spatio-temporal dynamics. Even though growth is localized within a layer of width *δ* at the colony frontier, the velocity field given by Eq. 2 will in general be non-zero in the interior of the colony. However, the velocity modulus decreases sharply with the distance from the frontier and is negligible outside the growth layer (Fig. 3D), so that flow in the colony occurs mostly within the growth layer.

**Fig. 3.**
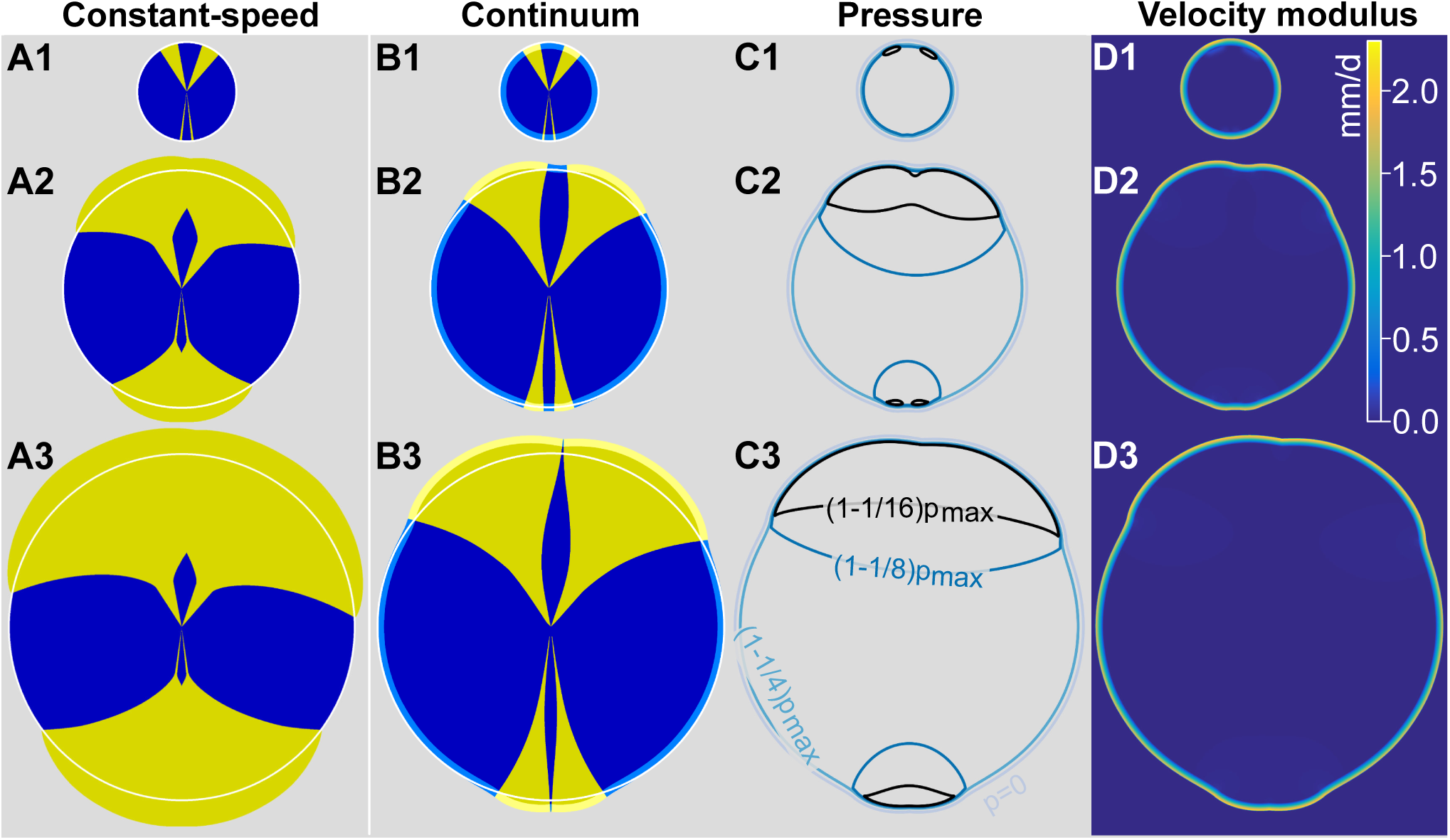
The constant-speed model inspired by geometrical optics (A1-A3) and the continuum model (B1-B3) make different predictions for the competition of yeast strains in growing colonies. Both models were integrated numerically, starting from the same initial condition shown in panels A1 and B1. The constant-speed model (A1-A3) fails to reproduce the thin, persistent filaments observed experimentally (Fig. 1A-A2), but such filaments are seen for the blue strain in the continuum model (B1-B3). The opening angle of yellow sectors increases faster in the constant-speed model (A1-A3) than in the continuum model (B1-B3), consistent with the experimental observations in Fig. 4. White contours are circles fit to the blue strain outer profile far from yellow sectors; they highlight the displacement of the blue strain close to yellow sectors in the continuum model. The growth layer is shown in B1-B3 using light yellow and light blue colors. Panels C1-C3 show pressure contours *p =* 0 and (1 – 1/*n*)*p*_*max*_, where *n =* 4,8,16 and *p*_*max*_ was evaluated at each time step. Panels D1-D3 show the local velocity modulus in the continuum model simulations and highlight that the displacement of strains takes place mostly within the growth layer. Model parameters are 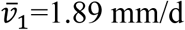, 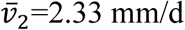 and *δ*=290 μm.

We find that simulations based on this continuum model can reproduce the experimental observation of the mechanical interaction of single-strain sectors within growing colonies (Fig. 3B), namely the displacement of the slower-growing strain in the proximity of a faster-growing sector and the prolonged survival and enhanced expansion speed of thin filaments of the slower-growing strain trapped between two sectors of the faster-growing one (Fig. 3B). Both these effects become more pronounced as the width, *δ*, of the growth layer increases. Note that the same observations are not predicted by a constant-speed model (5) inspired by geometrical optics in which each strain advances at a constant speed, perpendicular to the frontier of the colony (Fig. 3A). The constant-speed model is known to fail for the early dynamics of sweeping sectors (e.g., for sectors composed of cells with beneficial mutations), for example when the two walls bounding such sectors are close to each other (5). The temporal dynamics of a sweeping sector can be quantified by its opening angle, which is defined as the angle formed by the two lines connecting the center of the inoculum to the boundary between the two strains at the frontier of the colony (e.g., the angle *ϕ* in Fig. 1A). In the constant-speed model, the sector boundaries form logarithmic spirals, with their opening angle increasing logarithmically with the radius of the colony (5). In simulations of the continuum model, however, the dynamics of opening angles can be separated into two stages, which are best seen on a log-linear scale (continuous lines in Fig. 4D). Initially, opening angles increase exponentially with the radius *r*, defined at the point in which the two strains meet at the frontier of the colony (Fig. 4A, inset). At later times (or, equivalently, larger radii *r*), the rate of increase of sweeping sector opening angles versus *r* eventually tends to the constant-speed model prediction (Fig. 4D and Fig. S9), that is, opening angles increase logarithmically with *r*. We find that the continuum model results tend to the constant-speed model ones as the width of the growth layer *δ* tends to zero, while keeping the constant asymptotic expansion speed 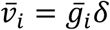 of each strain *i* constant by varying 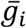 (Fig. S9). This observation makes sense because the constant-speed model, which is inspired by geometrical optics, assumes that each point at the colony frontier (corresponding to an extremely narrow growth layer) is the source of a circular growth wave in the infinitesimal time interval *dt*, and that the outline of the colony at time *t* + *dt* is given by the envelope of such waves (5).

**Fig. 4.**
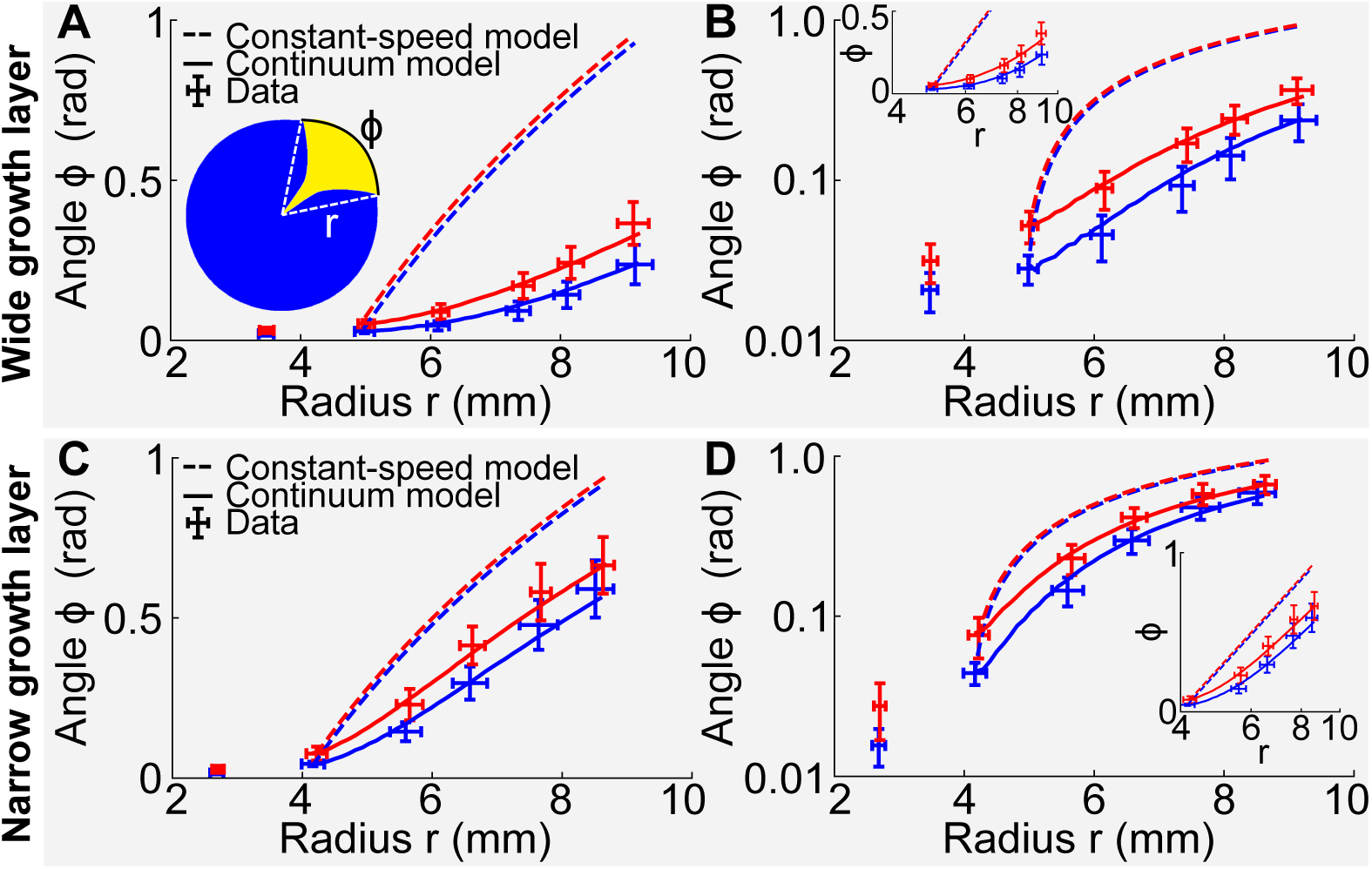
Dynamics of sector opening angles (data points are mean±SD) of strain yAG2 opening into strain yAG1 at 30°C after the inoculation of mixed-strain colonies with yAG2 relative frequency of 1%. The inset in (A) is a sketch of a sweeping sector within a colony. The radius *r* is defined as the distance from the center of the colony to the point at which the two strains meet at the frontier of the colony (dashed, white lines). The sector opening angle *ϕ* is the angle formed by the dashed, white lines, subtended by the arc shown in black. Experimental data were divided in two classes with equal numbers of sectors, based on the opening angle at the second measurement point (blue: small angle, red: large angle, SI Methods). Dashed curves are predictions of the constant-speed model. Continuous curves are best-fits to the continuum model. (A) Sector dynamics (red: 27 sectors, blue: 27 sectors) of the fitter strain in colonies with a wide layer of active growth at the frontier. (B) Same as panel A in log-linear scale. Inset: same data (with radii≥4 mm) in linear-log scale, where the constant-speed model prediction is a straight line. (C-D) Same as (A-B) for sector dynamics in colonies with a narrow layer of active growth at the frontier (red: 10 sectors, blue: 10 sectors). Experimental data show that opening angles increase more rapidly for narrow growth layers (C-D) than for wider ones (A-B). The transition between the exponential and logarithmic growth of opening angles with the radius *r* is clearly visible in (D). The constant-speed model (dashed lines) cannot reproduce the initial opening angles dynamics. The continuum model (continuous lines), on the other hand, can reproduce the data, using the growth layer width *δ* as a parameter. In (A-B), the best-fit parameter was *δ*=290 μm and in (C-D) it was *δ*=110 μm.

We tested whether our continuum model could reproduce the observed temporal evolution of the opening angles of sweeping sectors. We performed colony growth experiments at 30°C with the slower-growing strain yAG1 and the faster-growing strain yAG2 and measured the opening angles of sweeping yAG2 sectors that remained far from each other throughout the entire experiment (e.g., the four isolated yellow sectors at 2, 8, 9 and 11 o’ clock in Fig. 1A). Opening angles measured at different times and colony radii are shown in Fig. 4A-B and Fig. S10. We grouped the experimentally-measured opening angle trajectories of sweeping yAG2 sectors into two classes (small and large angles), based on the opening angle of each sector at the second measurement point, which was taken as the initial condition for the comparison with the models (see SI section 7 for an explanation). A log-linear plot highlights the initial exponential dependence of opening angles on the distance from the center of the colony to the points at which yAG1 and yAG2 sectors meet at the frontier of the colony (Fig. 4B). The constant-speed model (dashed lines in Fig. 4A-B, based on each strain’s expansion velocity measured in single-strain colony growth experiments), strongly overestimates the initial increase in the opening angles of sweeping yAG2 sectors. We tested if our continuum model could reproduce the experiments by fitting the continuum model to the experimental opening angles of sweeping sectors. The expansion velocities 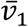 and 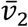 measured in single-strain colony growth experiments (SI section 8 and Fig. S11) constrained the products 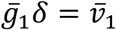 and 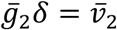 in the continuum model, assuming an identical growth layer width *δ* for the two strains. Thus, we could vary the width *δ* of the growth layer in the model (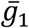 and 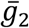 were then uniquely determined) and use it as a parameter to fit the sector opening angles. The resulting model fit is shown in Fig. 4A-B (continuous lines). The best-fit estimate is *δ*=290 μm (68% CI [260, 350] μm). We experimentally measured the characteristic length-scale over which growth affects the visible surface of yeast colonies and found it to be equal to *δ*_*exp*_ =530±120 μm (mean±SD, see SI section 9 and Fig. S12). The best-fit estimate of *δ* is smaller than *δ*_*exp*_, but this is not surprising given that the colony height goes to zero at the colony frontier, whereas the model assumes constant height everywhere (geometrical considerations suggest that *δ*_*exp*_ =2*δ*, see SI section 9).

The continuum model predicts that the opening angles of sweeping sectors increase faster for smaller values of *δ*. To test this prediction, we reduced the value of the growth layer width *δ* by reducing the wetness of the agar plates and measured the dynamics of the opening angles of sweeping sectors in this condition (Fig. 4C-D). We experimentally verified that the growth layer width was reduced, compared to wetter plates (SI section 9 and Fig. S12). In the experiment with a narrower growth layer width, even though the ratio of the expansion velocities of the two single strains 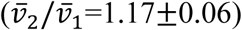 was slightly smaller than in the experiment with wider growth layer width 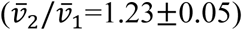, the opening angles increased more rapidly, consistent with the smaller growth layer width *δ*, thus verifying the continuum model’s prediction. Although the constant-speed model fails to predict correctly the early time dynamics of opening angles, it does a better job than for the previous experiment, as expected given the smaller value of *δ* and our observation that the continuum model tends to the constant-speed model in the limit *δ* → 0. As in the previous experiment, our continuum model can reproduce the experimental dynamics of sweeping sector opening angles by fitting the growth layer width to the data (Fig. 4C-D, continuous lines). The best-fit parameter for this experiment is *δ*=110±20 μm. In this case, one can observe the transition from the exponential (for small *ϕ* and small *r*) to logarithmic dependence of *ϕ* on *r* predicted by our continuum model (Fig. 4C-D). Such a transition is not discernible (or barely so) in the experiment described in the previous paragraph (Fig. 4A-B), due to the larger value of the growth layer width *δ* for wetter plates. In both experiments, the ratio 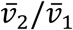 matches the ratio of experimentally-measured growth rates (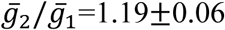, mean±SE), as expected based on the model prediction for the asymptotic expansion speed 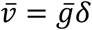.

## Discussion

Our experiments show that physical (pressure-induced) interactions between yeast strains within growing colonies reduce the efficiency of natural selection both by allowing less fit strains to persist and by slowing down the expansion of fitter strains. Slower-growing strains expand faster in parts of the colony that are close to faster-growing strains. When a slower-growing strain is trapped in a thin gap between two sectors of the faster-growing one, it expands at almost the same speed as the latter and forms a persistent, thin filament that can survive for very long times. A geometrical, toy model which neglects the contribution of curvature to the front propagation velocity (SI section 10) suggests that the thickness of these filaments decreases exponentially over a typical length scale 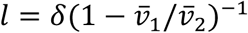, which increases with increasing growth layer width *δ* and the ratio 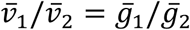. In the experiments, we found that thin filaments can persist longer than initially-larger sectors of the slower-growing strain, which are usually pinched off by sectors of the faster-growing strain that meet each other at the frontier of the colony subtending a substantial angle (SI section 11 and Fig. S15). The prolonged survival of thin, persistent filaments is favored by the small fluctuations of the filament boundaries relative to the filament thickness (SI section 12 and Fig. S16). Physical interactions also slow the initial dynamics of selective sweeps compared to the expectation of the constant-speed model, which assumes that the strains expand independently of each other except that two strains cannot occupy the same position. The magnitude of these effects increases with the width *δ* of the actively-growing layer at the frontier of the expanding colony. Our continuum model gives us an intuition for the physical origin of the experimental observations. Thin, persistent filaments are pushed towards the exterior of the colony by the surrounding faster-growing strain and benefit from the pressure field generated by the latter. Conversely, sweeping sectors of the faster-growing strain that have small initial opening angles are too small to generate enough pressure and thus initially expand more slowly than the predictions of the constant-speed model. Even though our continuum model is deterministic and thus cannot reproduce the stochastic extinction of small sectors of the faster-growing strain, which are sometimes observed in the experiments (see, e.g., Fig. 1A), it helps us to understand why such extinctions can occur. Sectors of the faster-growing strain initially expand at approximately the same speed as the surrounding slower-growing strain, making the competition between the two strains almost neutral, somewhat enhancing the importance of stochastic effects. Our experiments in time-varying environments show that the existence of thin, persistent filaments can play an important role in determining the outcome of competition between strains. In environments that fluctuate between two environmental states in which each strain has a selective advantage over the other one, the persistent filaments facilitate survival during detrimental environmental states and allow immediate recovery once the relative advantage of the two strains is reversed.

Because the reduced efficiency of natural selection described here is due to physical forces acting within dense cell populations, we believe such effects are relevant for a broad range of biological systems. We expect that the observed reduction in the effective selective advantage of a fitter strain (when sector widths are small) should be relevant for beneficial mutations that arise in a colony and manage to establish a clonal sector. In the initial phase of their dynamics, such sectors will expand at approximately the same rate as the surrounding wild type cells, and the expansion speed of the beneficial mutation will increase as its sector grows wider. This size-dependent selective advantage of a fitter strain or beneficial mutation is reminiscent of positive frequency-dependent selection (24), in which the fitness of a genotype increases with its frequency in the population. In the context of our work, the frequency dependence is caused by the pressure field induced by cell division. Experimental (25) and theoretical (26) studies have shown that populations undergoing range expansion accumulate deleterious mutations as a result of consecutive founder events and genetic drift being stronger at the frontier in spatial (compared to well-mixed) populations, due to their reduced effective population sizes. Our work suggests that deleterious mutations have higher chances of being maintained in dense, expanding microbial populations than expected based on genetic drift alone, because the effects of differences in fitness are reduced by physical interactions. Our findings are relevant for the fate of drug-resistant mutations that arise in microbial colonies or biofilms. Such mutations are typically associated with fitness deficits. Our work suggests that drug-resistant mutants (e.g., antibiotic-resistant bacteria) emerging in the outer layer of an expanding colony may benefit from the physical interaction with the surrounding wild-type cells and survive for long times in the actively-dividing frontier of the colony in the absence of the drugs that they confer resistance to. The application of a drug would reverse the relative fitness of drug-resistant mutants and wild-type cells and allow the drug-resistant mutant to take over the population, similar to our experiments where temperature varies in time for temperature-sensitive strains. We thus expect that the physical interactions between different genotypes investigated here would favor the maintenance of genetic diversity within dense microbial populations and allow such populations to withstand time-dependent fluctuations of the environment or the application of antimicrobial drugs. Finally, our findings may extend beyond microbial populations and inform us about the dynamics of pre-vascular tumor growth (7, 27). Within tumors, various cell lineages compete for space and form spatially-coherent clusters that are reminiscent of the genetic de-mixing found in colony-growth experiments (28). A recent study (28) has shown that individual tumors can have high levels of genetic diversity and argued that such high-diversity levels can only be explained by non-Darwinian selection, i.e., neutral competition between different clones. The reduced efficiency of natural selection highlighted here, transposed to the three-dimensional environment at the frontier of a spherical cell cluster, is one possible mechanism to explain the apparent neutrality of inter-clone competition within such tumors.

## Methods

### Strains

The genotypes of all strains used in this study are reported in Table S1. In fluorescent images, yAG1 is portrayed in blue, yAG2 in yellow, yAG19 in cyan and yAG20 in magenta. Strains’ growth rates were measured by imaging time-lapse videos of micro-colonies of each strain on standard YPD agar plates at 30°C using an inverted microscope and counting the number of cells in each micro-colony at successive time points. The ratio of experimentally-measured growth rates 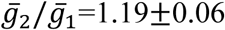 (mean±SE) was estimated by randomly sampling micro-colonies of the two strains, computing their growth rate ratio and taking the mean across 10^6^random samples.

### Colony-growth experiments

Colony-growth experiments were conducted on 1% agarose plates with YPD (Yeast extract-Peptone-Dextrose) medium at 30°C, unless otherwise stated. Plates were poured two days before inoculation and stored at room temperature. Strains were pre-grown at 30°C overnight in liquid YPD. Before inoculation on agarose plates, 200 μL of each overnight culture were spun down for 1 min at 9400 *g* and re-suspended in 1 mL of 1% PBS (Phosphate-Buffered saline). The densities of such suspensions were measured with a Coulter Counter and brought to a density of 3·10^7^cells/mL by adding 1% PBS or removing the supernatant in appropriate amounts. If required, suspensions were mixed in the desired ratios. A 0.6 μL droplet was pipetted on each agarose plate.

### Sweeping sectors dynamics experiment

We performed two sets of experiments to measure the dynamics of sweeping yAG2 sectors at different growth layer widths at 30°C. For the wide growth layer experiment, we poured 7 mL YPD 1% agarose medium into 35 mm Petri dishes. After one day from pouring, solidified YPD agarose discs were transferred to 100 mm Petri dishes, three discs for each larger dish. One droplet of either a single-strain or a mixed-strain suspension (yAG1 and yAG2 were at a 99%-1% frequency) was inoculated on the surface of each disc. 7 mL of liquid YPD medium were added to the Petri dish and substituted daily to ensure that nutrients did not become depleted. Colonies were imaged twice per day with a Zeiss Lumar stereoscope. For the narrow growth layer, we poured 33 mL YPD 1% agarose medium into 100 mm Petri dishes. One droplet of either a single-strain or a mixed-strain suspension (yAG1 and yAG2 were at a 99%-1% frequency) was inoculated on the surface of each plate. Colonies were imaged once per day with a Zeiss Lumar stereoscope.

### Radial expansion velocities

Single-strain colony radii were measured by fitting circles to each single-strain colony bright-field image. Radial expansion velocities were computed by fitting the radius-vs-time data to straight lines, separately for each colony. The mean radial expansion velocity of each strain was computed as the mean slope of the radius-vs-time fits. For the wide growth layer experiment, yAG1 and yAG2 expansion velocities were measured across 10 and 9 single-strain colonies, respectively. The mean radii at different times are shown in Fig. S11A. The measured velocities were 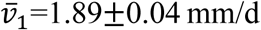 (mean±SD) and 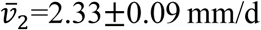 (mean±SD). For the narrow growth layer experiment, yAG1 and yAG2 expansion velocities were measured across 11 single-strain colonies for each strain. The mean radii at different times are shown in Fig. S11C. The measured velocities were 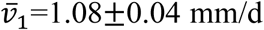 (mean±SD).and 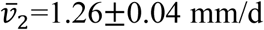 (mean±SD).

### *cdc26*Δ *mutant experiments*

We performed colony-growth experiments with the strains yAG19 and yAG20 to show that filaments and strain displacements can occur without proliferation of the less-fit strain. These experiments were done with 100 mm Petri dishes filled with 33 ml of 1% agarose YPD medium. We grew single-strain and mixed-strain colonies of yAG19 and yAG20 (yAG19 and yAG20 were at a 5%-95% frequency) for two days at 28°C and then moved them to the stage of a stereoscope incubated at 37°C. Colonies were imaged every 10 min for 23 h. The same experiment was performed with yAG2 and yAG20 single-strain and mixed-strain colonies and the results do not differ qualitatively.

### Time-varying temperature experiments

We performed time-varying temperature experiments with 100 mm Petri dishes filled with 33 mL of 1% agarose YPD medium. We inoculated 11 plates with a mixed-strain 0.6 μL droplet (yAG1 and yAG2 were at a 50%-50% frequency). Three of these plates were incubated at 30°C for 2 days, then at 12°C for 9 days and then at 30°C for 14 days (Fig. S3). Three plates were incubated at 30°C for 3 days, then at 12°C for 16 days and then at 30°C for 10 days (Fig. 2A-C and Fig. S4). Three plates were incubated at 30°C for 3 days, then at 12°C for 20 days and then at 30°C for 9 days (Fig. S5). One plate was incubated at 30°C for 13 days and then at 12°C for 18 days (Fig. 2D-L). We inoculated 6 plates with a 0.6 μL droplet of yAG1 and other 6 with yAG2. Three of each strain’s plates were stored at room temperature overnight and then moved to a 12°C incubator for the rest of the experiment. The remaining three plates for each strain were placed in a 30°C incubator for the entire experiment. We took frequent fluorescent and bright-field images of all plates. Single-strain colonies were used to measure single-strain radial expansion velocities.

### Continuum and constant-speed models

The continuum and the constant-speed models were solved numerically using level-set methods (29) and isotropic discrete operators from lattice hydrodynamics (30). Details on the numerical implementation are in the SI Methods.

## Acknowledgements

AG thanks Sauro Succi for helpful discussions on the numerical integration of the continuum model and the Harvard Center for Biological Imaging and Douglas Richardson for helping with the growth rates measurement on agar plates. We thank the members of the AWM and DRN groups for valuable suggestions. AG was supported by research fellowships from the Swiss National Science Foundation, projects P2ELP2_168498 and P400PB_180823, and work on this project was supported by the Human Frontier Science Program Grant RGP0041/2014 to AWM and DRN. Work by AG and DRN was supported by the National Science Foundation, through grants DMR1608501 and via the Harvard Materials Science Research and Engineering Center via grant DMR1435999.

**Author contributions**
A.G., D.R.N., A.W.M. designed the research and wrote the paper, A.G. performed the research and analyzed the data.

